## Supplementary material for "The E3 ubiquitin ligase RNF220 maintains hindbrain *Hox* expression patterns through regulation of WDR5 stability": Figure supplements information

Huishan Wang et al.

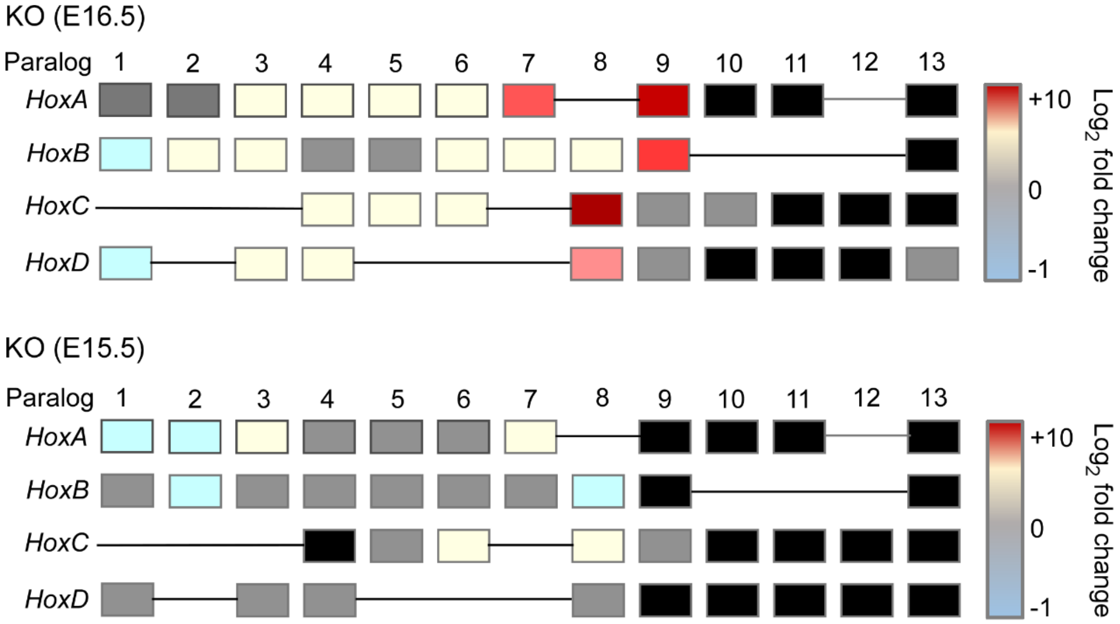


**Figure 1-figure supplement 1.** *Hox* genes exhibited de-repression in hindbrain of *Rnf220*$^{-/-}$ embryos at late developmental stages. Heatmap of *Hox* expression in hindbrain of *Rnf220*$^{-/-}$ mouse embryos at E16.5 and E15.5. Expression of each *Hox* gene was analyzed by qRT-PCR against GAPDH. Each *Hox* gene in wild-type controls was set to 1 (n=2 mice per group). Blank color represents the expression of that *Hox* is low and exceed detection range. KO, knockout.

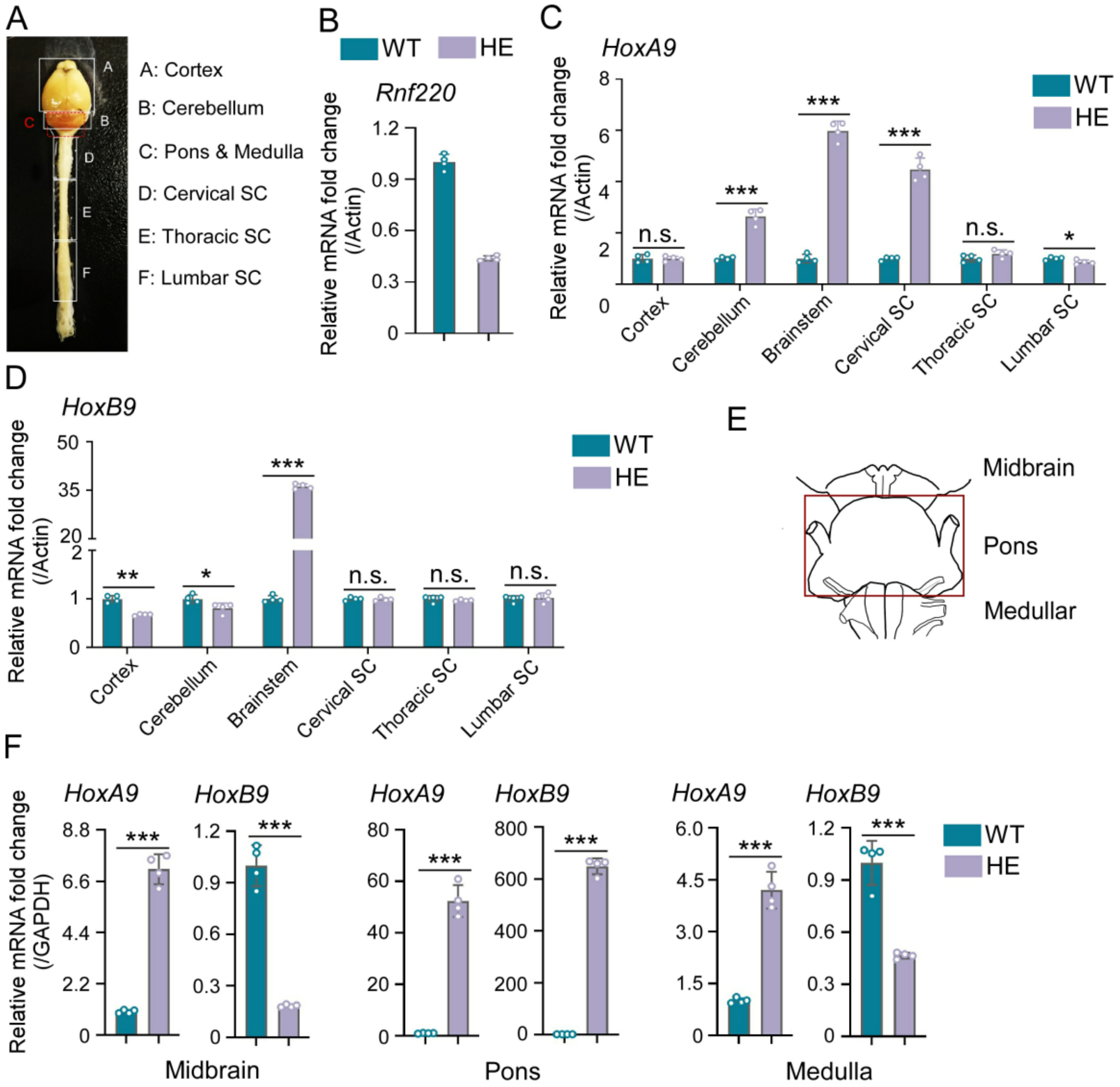


**Figure 1-figure supplement 2.** *Hox* genes were up-regulated in brainstem of *Rnf220*$^{+/-}$ mice. (**A**) Diagram of central neural system (CNS) in mice. (**B**) qRT-PCR analysis of *Rnf220* expression in CNS of *Rnf220*$^{+/-}$ and control mice. (**C-D**) qRT-PCR analysis of *HoxA9* (C) and *HoxB9* (D) expression in each CNS section of *Rnf220*$^{+/-}$ and control mouse (n=2 mice per group). (**E**) Diagram of mouse brain stem. (**F**) qRT-PCR analysis of mRNA levels of *HoxA9* and *HoxB9* in indicated sections of WT and *Rnf220*$^{+/-}$ mouse brainstems (n=2 mice per group). WT, wild-type. HE, heterozygote. SC, spinal cord. n.s., not significant. *$p < 0.05$, **$p < 0.01$, ***$p < 0.001$.

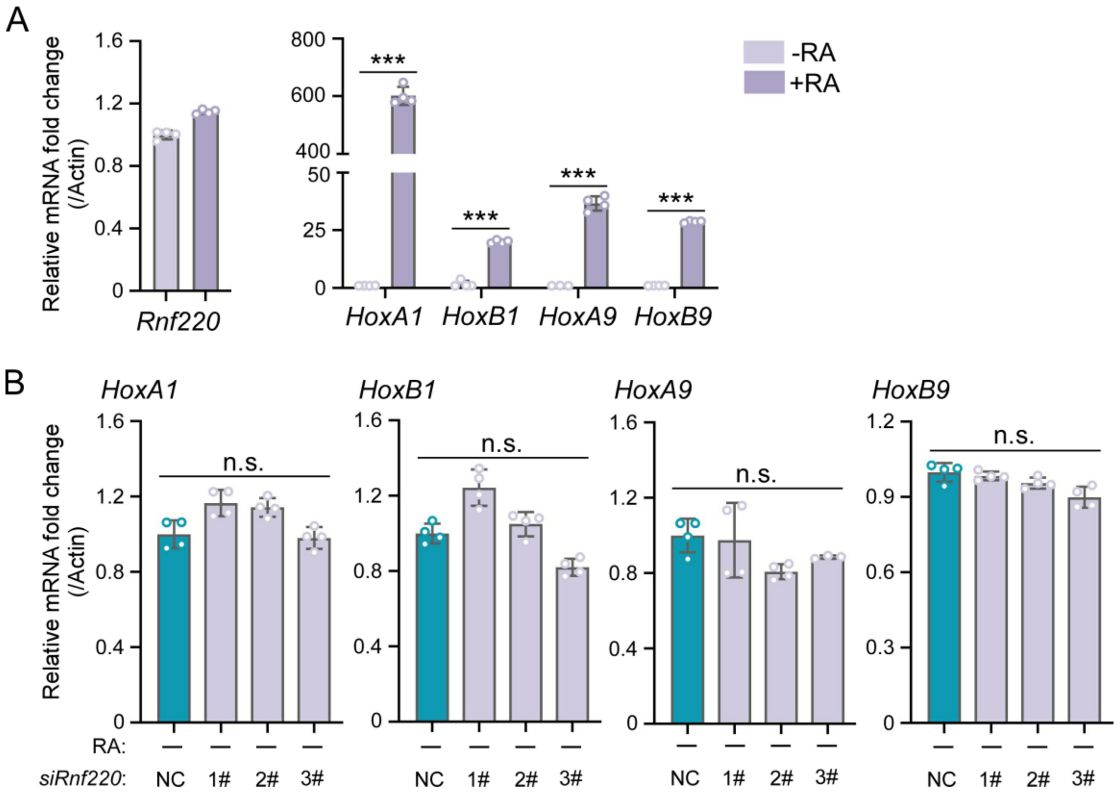


**Figure 1-figure supplement 3.** Expression levels of *Hox* genes were not affected by *Rnf220* knockdown in P19 cell line without RA induction. (**A**) qRT-PCR analysis of *Rnf220* and indicated *Hox* genes expression after RA treatment in P19 cells. (**B**) qRT-PCR analysis the expression levels of *HoxA1*, *HoxB1*, *HoxA9*, and *HoxB9* in *Rnf220* knockdown P19 cells without RA induction. RA: retinoic acid. n.s., not significant. ***$p < 0.001$.

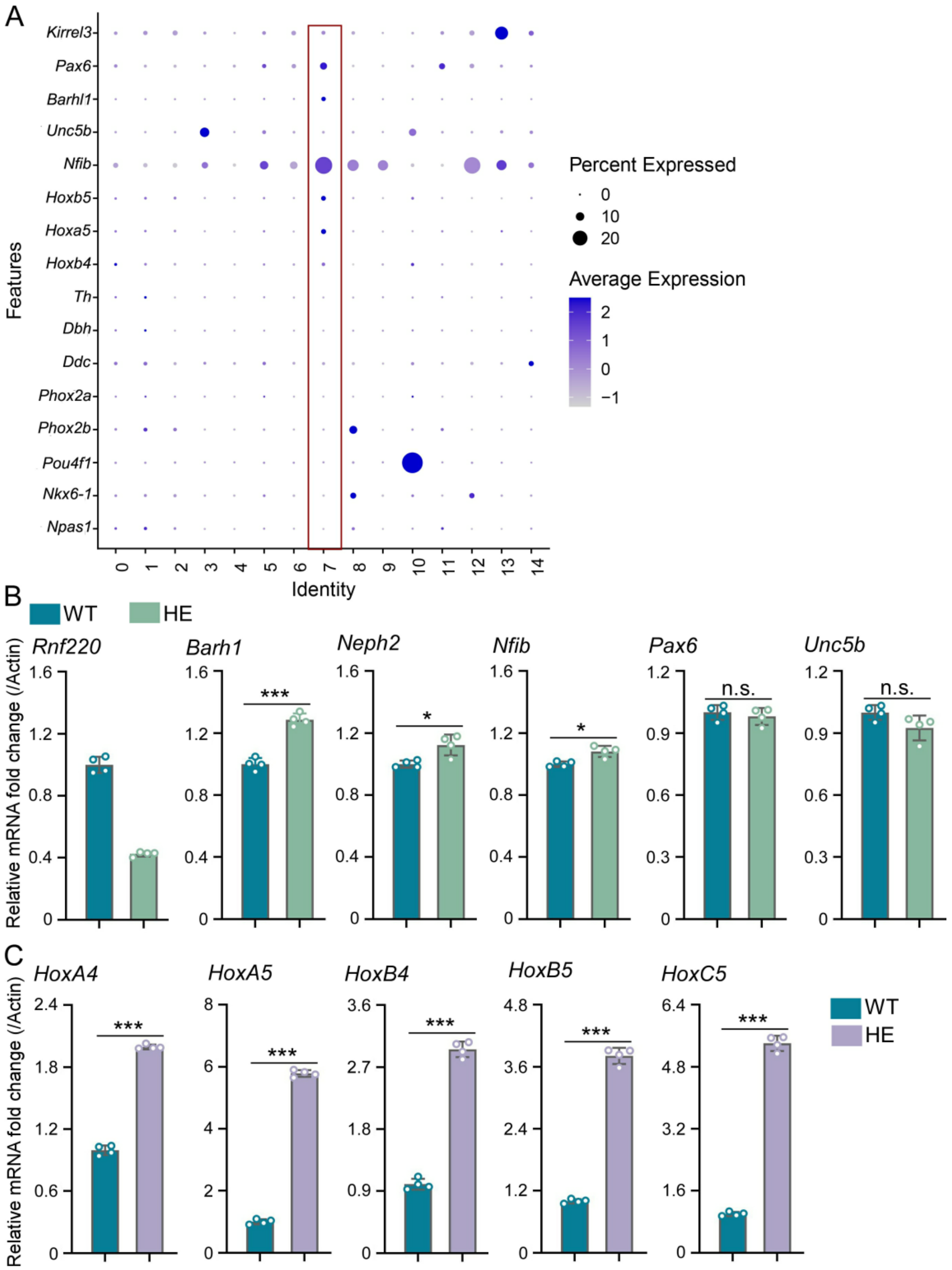


**Figure 2-figure supplement 1.** *Hox* genes were up-regulated in PN of *Rnf220*$^{+/-}$ mice. (**A**) Heatmap of snRNA-seq data showing expression levels of PN markers (*Pax6, Barhl1, Unc5b*, and *Nfib*) among 15 identified cell clusters (n= 3 mice per group). (**B-C**) qRT-PCR analysis of expression levels of indicated PN markers (B) and *Hox* genes (C) in pons of *Rnf220*$^{+/-}$ and WT mice (n= 3 mice per group). WT, wild-type. HE, heterozygote. n.s., not significant. *$p$ < 0.05, ***$p$ < 0.001.

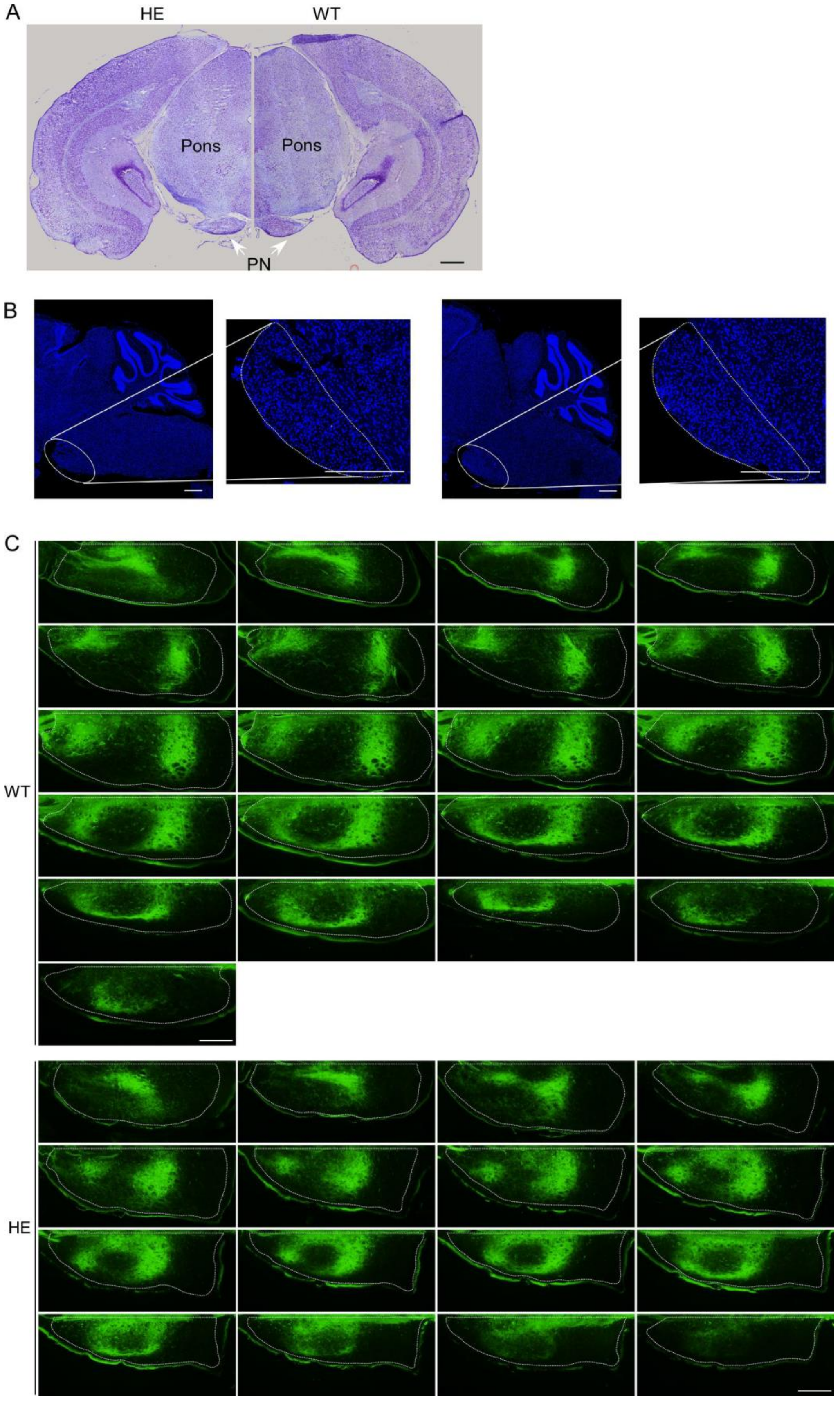


**Figure 2-figure supplement 2.** PN showed no structure difference but disorganized projection pattern from motor cortex between WT and *Rnf220*$^{+/-}$ mice. (**A**) Nissl staining showed the pons structure in adult (2 months) WT and *Rnf220*$^{+/-}$ mice. Scale bars, 500 μm. (**B**) DAPI labeling PN structure in adult (2 months) WT and *Rnf220*$^{+/-}$ mice. Scale bars, 500 μm. (**C**) Continuous slice of projection from motor cortex to PN in adult (2 months) WT and *Rnf220*$^{+/-}$ mice. Scale bars, 200 μm. WT, wild-type. HE, heterozygote. PN, pontine nuclei.

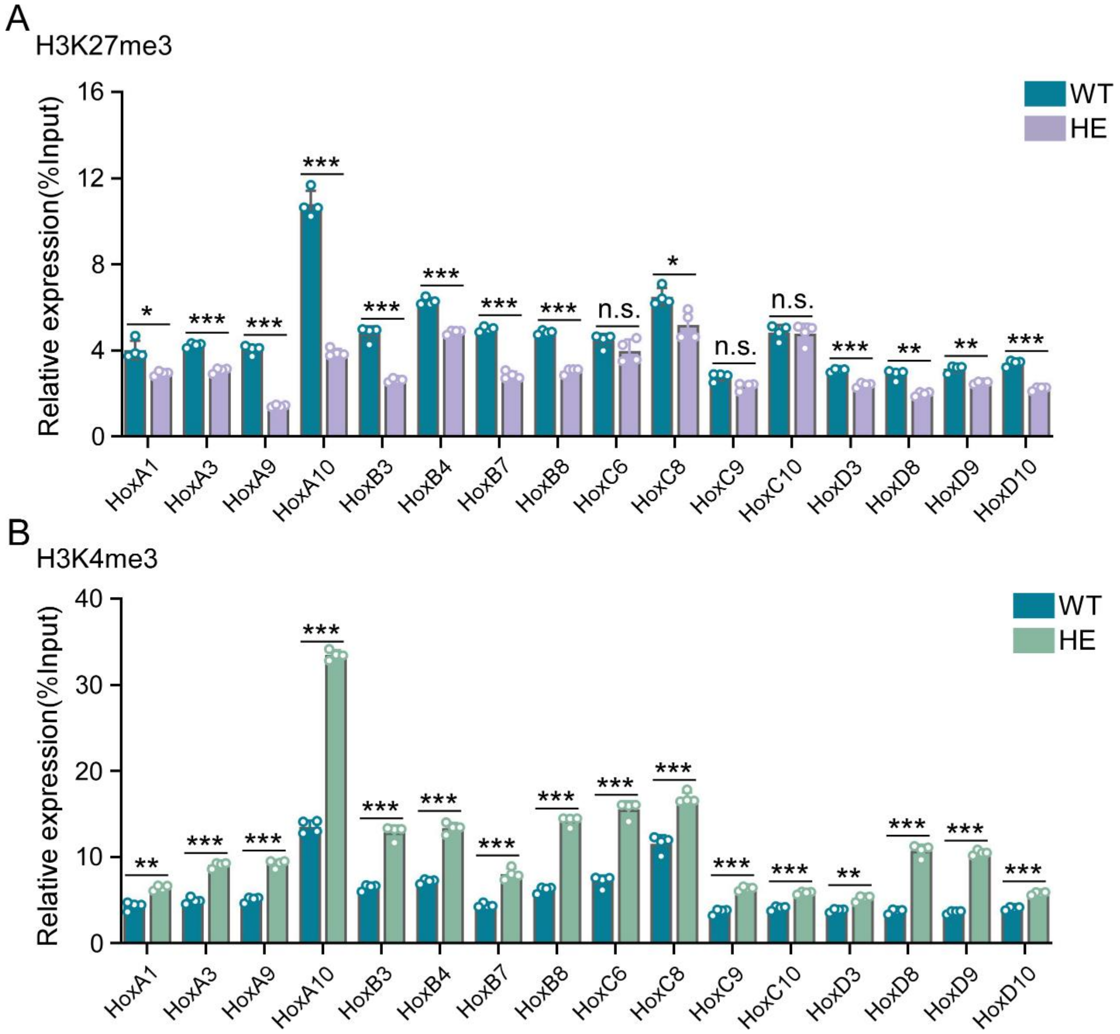


**Figure 3-figure supplement1.** Repressive epigenetic modification was down-regulated while activated epigenetic modification was up-regulated in promoter regions of indicated *Hox* genes in hindbrains of *Rnf220*$^{+/-}$ mice. **(A-B)** ChIP-qRT-PCR analysis of repressive epigenetic modification (H3K27me3) (A) and activated epigenetic modification (H3K4me3) (B) levels in promoter regions of indicated *Hox* genes in hindbrains of *Rnf220*$^{+/-}$ and WT mice (n=2 mice per group). WT, wild-type. HE, heterozygote. n.s., not significant. *$p$< 0.05, **$p$ < 0.01, ***$p$ < 0.001.

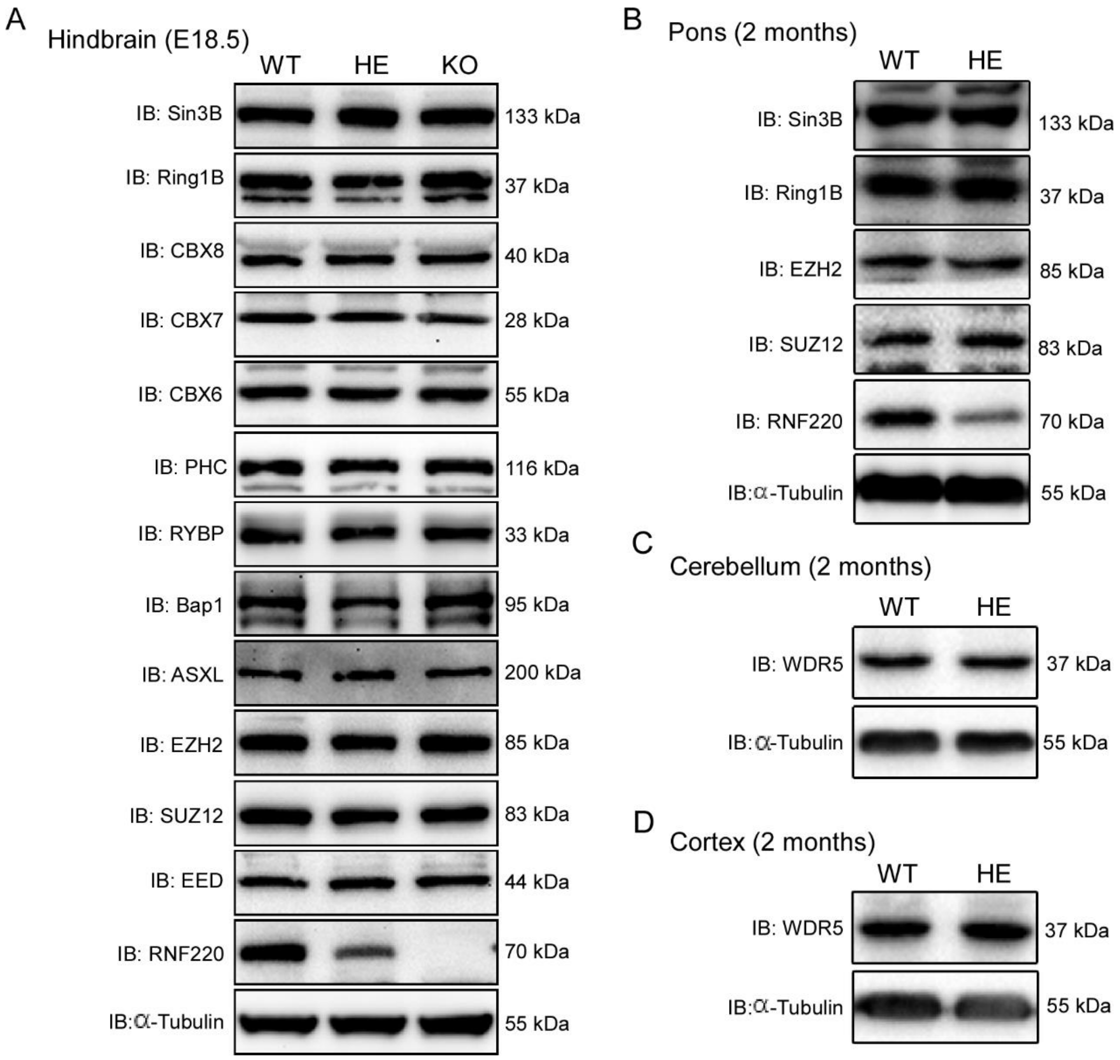


**Figure 3-figure supplement 2.** Protein levels of the indicated core components of PRC1 and PRC2 complex in indicated mouse brain tissues of different genotypes. (**A**) Western blot analysis of protein levels of core components of PRC1 and PRC2 complexes in hindbrain of WT, *Rnf220*$^{+/-}$, and *Rnf220*$^{-/-}$ mouse embryos at E18.5. (**B**) Western blot analysis of protein levels of core components of PRC1 and PRC2 complexes in pons of adult *Rnf220*$^{+/-}$ and WT mice. (**C, D**) Western blot analysis of protein levels of WDR5 in cerebellum and cortex of adult *Rnf220* $^{+/-}$ and WT mice. WT, wild-type; HE, heterozygote; KO, knockout; IB, immunoblot.

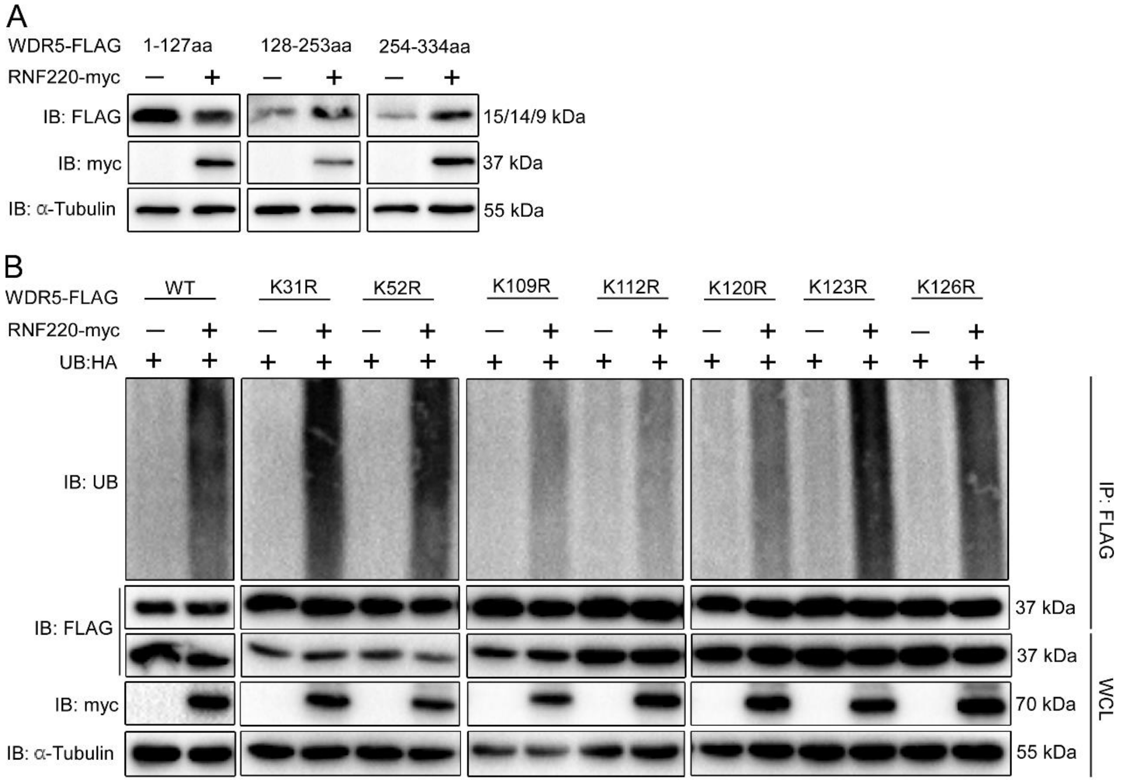


**Figure 4-figure supplement 1.** RNF220 interacted with and targeted WDR5 for polyubiquitination at multiple lysine sites. (**A**) Western blot analysis the levels of three WDR5 truncated proteins in HEK293 cells when co-transfected with RNF220 or not. (**B**) *In vivo* ubiquitination analysis of ubiquitination status of indicated WDR5 KR mutants when co-expressed with RNF220 or not in HEK293 cells. IB, immunoblot. IP, immunoprecipitation. WCL, whole cell lysate. K31R, K52R, K109R, K112R, K120R, K123R, or K126R, substitution of lysine with arginine in WDR5 at indicated positions.

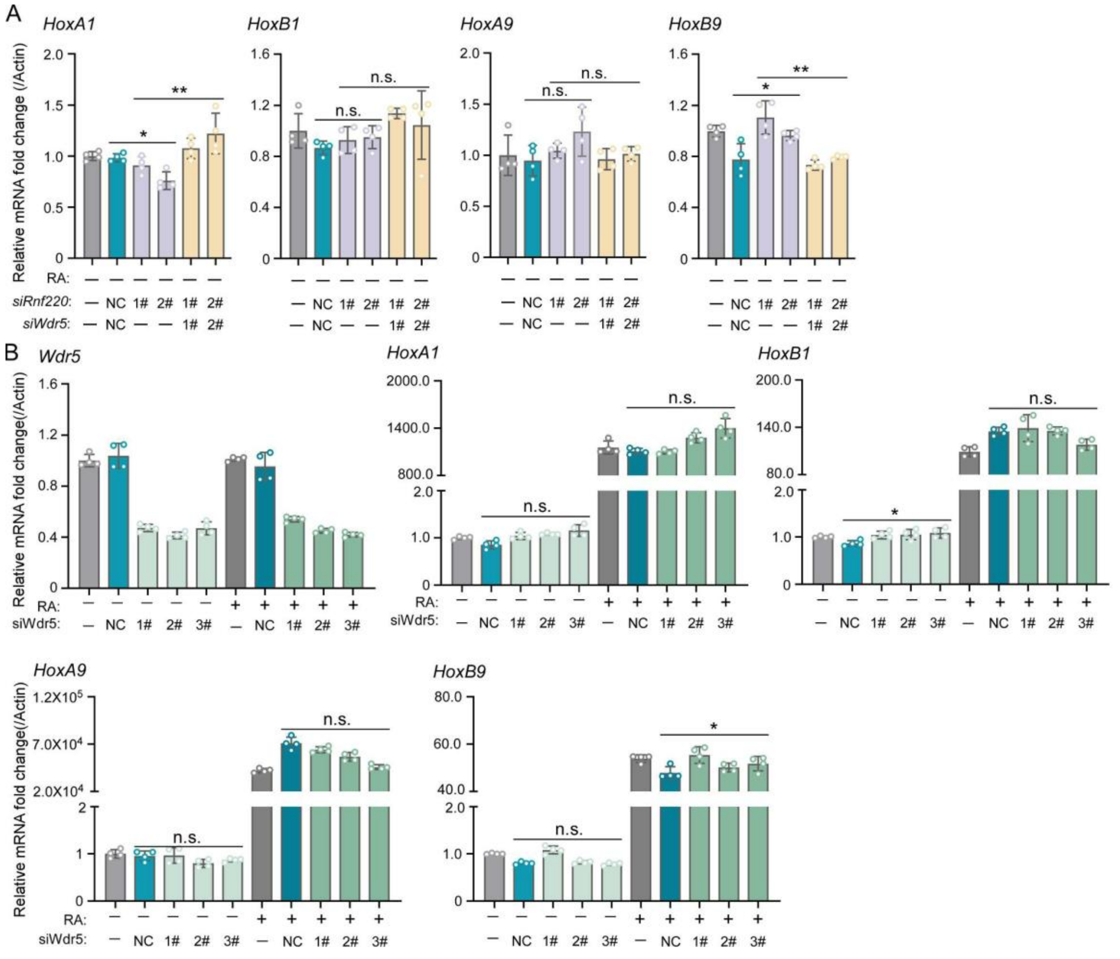


**Figure 5-figure supplement 1.** WDR5 knockdown or without RA treatment had no effect on *Hox* genes in P19 cell line. (**A**) qRT-PCR analysis showing the expression levels of *HoxA1*, *HoxB1*, *HoxA9*, *HoxB9* when transfected siRnf220 or both siRnf220 and siWdr5 without RA treatment. (**B**) qRT-PCR analysis of mRNA levels of *Wdr5*, *HoxA1*, *HoxB1*, *HoxA9*, and *HoxB9* when *Wdr5* was knocked down by siRNA transfection in P19 cells with or without RA treatment. Bar graphs (mean ± SD) show the relative levels normalized against control group without siRNA or RA treatment. RA: retinoic acid. n.s., not significant. *$p < 0.05$.

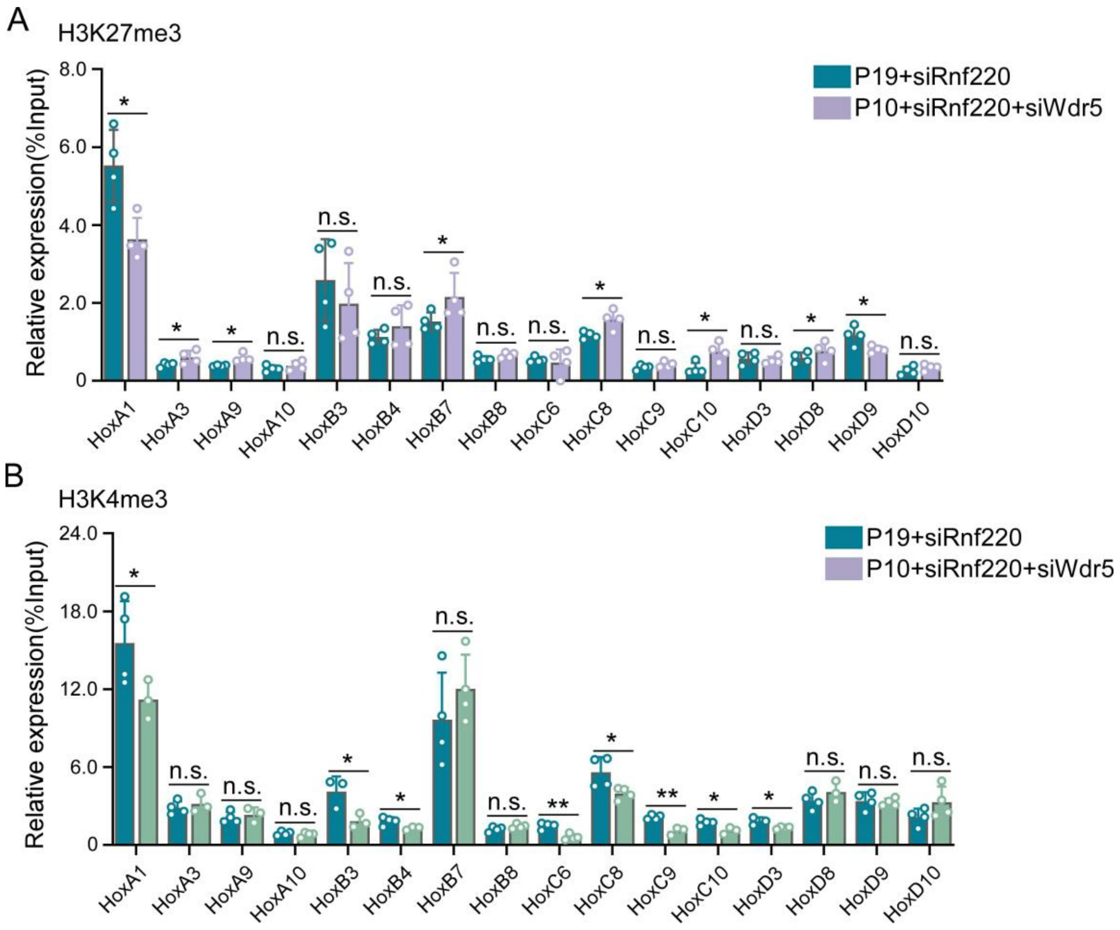


**Figure 6-figure supplement 1. *Rnf220* and *Wdr5* co-suppression recovered *Hox* epigenetic modification to a certain degree.**

(**A-B**) ChIP-qRT-PCR analysis of repressive epigenetic modification (H3K27me3) (A) and activated epigenetic modification (H3K4me3) (B) levels in promoter regions of indicated *Hox* genes in P19 cell line transfected with siRnf220 or both siRnf220 and siWdr5. n.s., not significant. *$p$< 0.05, **$p$ < 0.01.
